# A systematic analysis of the qualitative and quantitative evidence on the purpose of animal-assisted therapy in improving patients wellbeing

**DOI:** 10.1101/2023.08.10.552837

**Authors:** K. Himanshu, K. Gunjan, Riya Mukherjee, Ramendra Pati Pandey, Chung-Ming Chang

**Author notes:** Correspondence (C.M.C); (R.P.P).

## Abstract

**Background:** Animal-assisted therapy, also known as pet therapy, is a therapeutic intervention that involves animals to enhance the well-being of individuals across various populations and settings. This systematic study aims to assess the outcomes of animal-assisted therapy interventions and explore the associated policies.

**Methods:** A total of 16 papers published between 2015 and 2023 were selected for analysis. These papers were chosen based on their relevance to the research topic of animal-assisted therapy and their availability in scholarly databases. Thematic synthesis and meta-analysis were employed to synthesize the qualitative and quantitative data extracted from the selected papers.

**Results:** The analysis included sixteen studies that met the inclusion criteria and were deemed to be of moderate or higher quality. Among these studies, four demonstrated positive results for therapeutic mediation and one for supportive mediation in psychiatric disorders. Additionally, all studies showed positive outcomes for depression and neurological disorders. Regarding stress and anxiety, three studies indicated supportive mediation while two studies showed activating mediation.

**Conclusion:** The overall assessment of animal-assisted therapy shows promise as an effective intervention in promoting well-being among diverse populations. Further research and the establishment of standardized outcome assessment measures and comprehensive policies are essential for advancing the field and maximizing the benefits of animal-assisted therapy.

## Introduction

The inclusion of animals in psychological treatment is not a recent or uncommon practice. Throughout history, there has been an understanding of the positive impact animals can have on human well-being[1]. This connection between humans and animals is deeply ingrained in our collective subconscious, influencing our emotional experiences. [2] The earliest documented instance dates back to the late 18th century, when animals were introduced into mental health institutions to enhance social interaction among patients[3, 4]. Today, numerous programs worldwide incorporate animals to varying degrees in their services. These programs are particularly beneficial for individuals who have experienced trauma, including those diagnosed with posttraumatic stress disorder (PTSD), schizophrenia, Alzheimer’s disease, Autism, etc[4, 5].

In the past 50 years, the field of human-animal interactions (HAI) and specifically, animal- assisted therapy (AAT), has made significant advancements and progress. AAT is a therapeutic approach that utilizes animals to improve overall health and well-being. It encompasses emotional, psychological, and physical interactions between individuals, animals, and the environment[6]. AAT interventions involve qualified treatment providers facilitating interactions between patients and animals with specific therapeutic goals in mind. These interventions often involve collaborative activities between human-animal teams, aiming to promote therapeutic and supportive outcomes[7]. AAT interventions contribute to individuals’ well-being, supporting physical health, and improving cognitive, emotional-affective, and social aspects, leading to enhanced emotional well-being, reduced anxiety, and decreased stress levels[8–10].

Research on therapies involving human-animal interaction has focused on specific animals such as: dogs, cats, or horses, as well as specific populations such as those with autism[11]. Dogs, in particular, are commonly preferred for therapy due to their exceptional bond with humans in modern times. Over thousands of years of shared evolutionary history[1], Dogs have acquired adept socialization skills with humans through processes of domestication and natural selection. They have become our loyal companions, developing unique social skills for interacting with humans.. For instance, studies indicate that dogs possess a sensitivity to our emotional states[12] and can interpret our social cues[13], even engaging in sophisticated communication through behaviors like gaze alternation[14]. Furthermore, dogs are capable of forming intricate attachment relationships with humans, resembling the bonds found in relationships between infants and caregivers[15]. suggest that among the various animals involved in animal-assisted therapy (AAT), dogs tend to exhibit superior interactions with people compared to other species, benefiting both children and adults[6].

This systematic review and meta-analysis specifically examine animal-assisted therapies, with an emphasis on both quantitative and qualitative approaches. These approaches involve incorporating human-animal interaction alongside, or in addition to, established and professional forms of various therapies. The special bonds formed between humans and animals are recognized as essential catalysts for transformation and are held in high regard, similar to the therapist-client relationship.

## 1. Material and method

### 2.1 Search strategy

The meta-analysis was carried out in accordance with the methodologies outlined in the esteemed Cochrane Handbook for Systematic Reviews of Interventions [16], and the findings were reported in compliance with the Preferred Reporting Items for Systematic Reviews and Meta-Analyses (PRISMA) guidelines[17]. To ensure comprehensive coverage, electronic databases were meticulously searched up till June 2023. A total of five English-language electronic databases, including PubMed, Web of Science, Clinical Trials, Science Direct, and Google Scholar, were meticulously explored. This thorough exploration entailed employing a combination of pertinent controlled vocabulary terms (such as MeSH) and relevant free text terms. The search strategy employed can be summarized as follows: (animal assisted therapy OR animal assisted intervention OR animal assisted activity OR animal activity interaction OR animal assisted method OR animal facilitated therapy OR pet therapy OR canine assisted therapy OR dog assisted therapy) AND (quasi-experimental study OR randomized controlled trial) AND (pain OR anxiety OR depression OR blood pressure OR BP OR heart rate OR HR) AND (“work-related stress” OR “workplace health” OR “employee well-being” OR “burnout”) AND (tumor OR malignant OR carcinoma OR oncology OR hospitalization OR hospitalized patients OR inpatients). By utilizing this extensive and refined approach, the meta-analysis aimed to capture a comprehensive body of evidence pertaining to the effects of animal-assisted interventions on various health outcomes.

### 2.2 Inclusion and exclusion criteria

The inclusion criteria were set based on the Patient, Intervention, Comparison, and Outcome (PICOS) framework: (i) Studies evaluating the effects of animal-assisted therapy, animal-assisted intervention, or animal-assisted activity. (ii) The effects of animal interactions on health and wellbeing (including depression, agitation, loneliness and stress and quality of life), social interaction, engagement, physical function, behavioral symptoms, medication use and adverse events. (iii) The articles should to be published in English (iv) Studies available in full-text format. (v) Studies utilizing quasi-experimental designs or randomized controlled trials. To maintain the rigor and relevance of the study, publications that lacked sufficient information regarding the therapy or did not involve an animal intervention were excluded from consideration.

### 2.3 Data synthesis

In our study, we employed thematic synthesis as a method to assess the eligibility and quality of the articles[18]. Each article was independently reviewed to determine its suitability for inclusion. We followed a traditional methodology for evaluating the papers, which involved examining factors such as the presence of adequate control group(s), control of confounders, randomization, well-described experimental design, and relevant outcome variables. Articles that met these criteria were selected and organized into a single sheet using Microsoft Office Excel® (2019). For the included studies, we extracted and compiled various data points into a structured table. This information encompassed the author’s name, country of publication, year of publication, patient characteristics (including sample size, age, gender, and target group), type of study, study design, description of AAT, type of intervention, control group details, study duration, outcomes measured, and the author’s conclusions. To effectively manage the papers, we utilized Mendeley software (version 1.19.8, Elsevier, London, UK).

### 2.4 Classification

In order to determine the specific contexts in which AATs are effective, we classified the interventions into three categories: supportive mediation, therapeutic mediation, and activating mediation. This categorization was based on the criteria outlined in Table 3.

## 2. Results

### 3.1 Study selection

The outcome of the search is depicted in Fig. 1.The search process resulted in 968 unique articles after initial searches from various electronic databases like PubMed, Web of Science, Science Direct, and Google Scholar, which yielded 942 articles. An additional 26 articles were extracted from other sources. After eliminating duplicate articles, the total number of articles was reduced to 507. Subsequently, the articles were assessed based on their title and abstract to determine eligibility. Among the initial pool, 389 articles were excluded as they did not meet the eligibility criteria, mainly due to the lack of relevance to AAT. After reading the full text of the remaining articles, 102 more articles were excluded. Out of these 102, 60 did not meet the inclusion criteria, and 42 were excluded due to being classified as Non-occupational mixed group or having Unrepresentative results. Finally, a total of 16 studies that met the inclusion criterion were included in the final analysis. The findings and details of these 16 studies are summarized in Tables 1 and 2.

**Figure 1:**
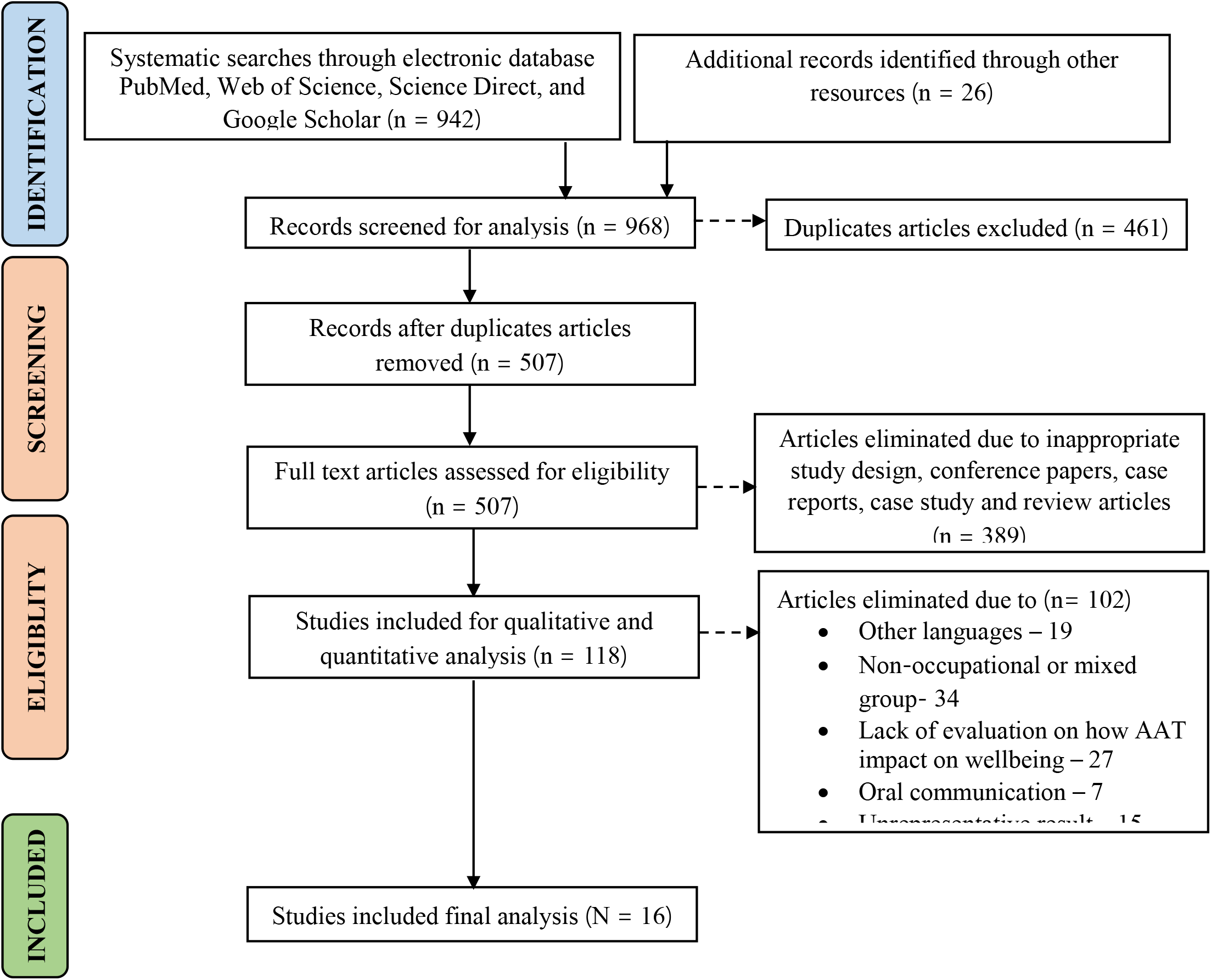
Literature screening flowchart (PRISMA)

**Table 1:**
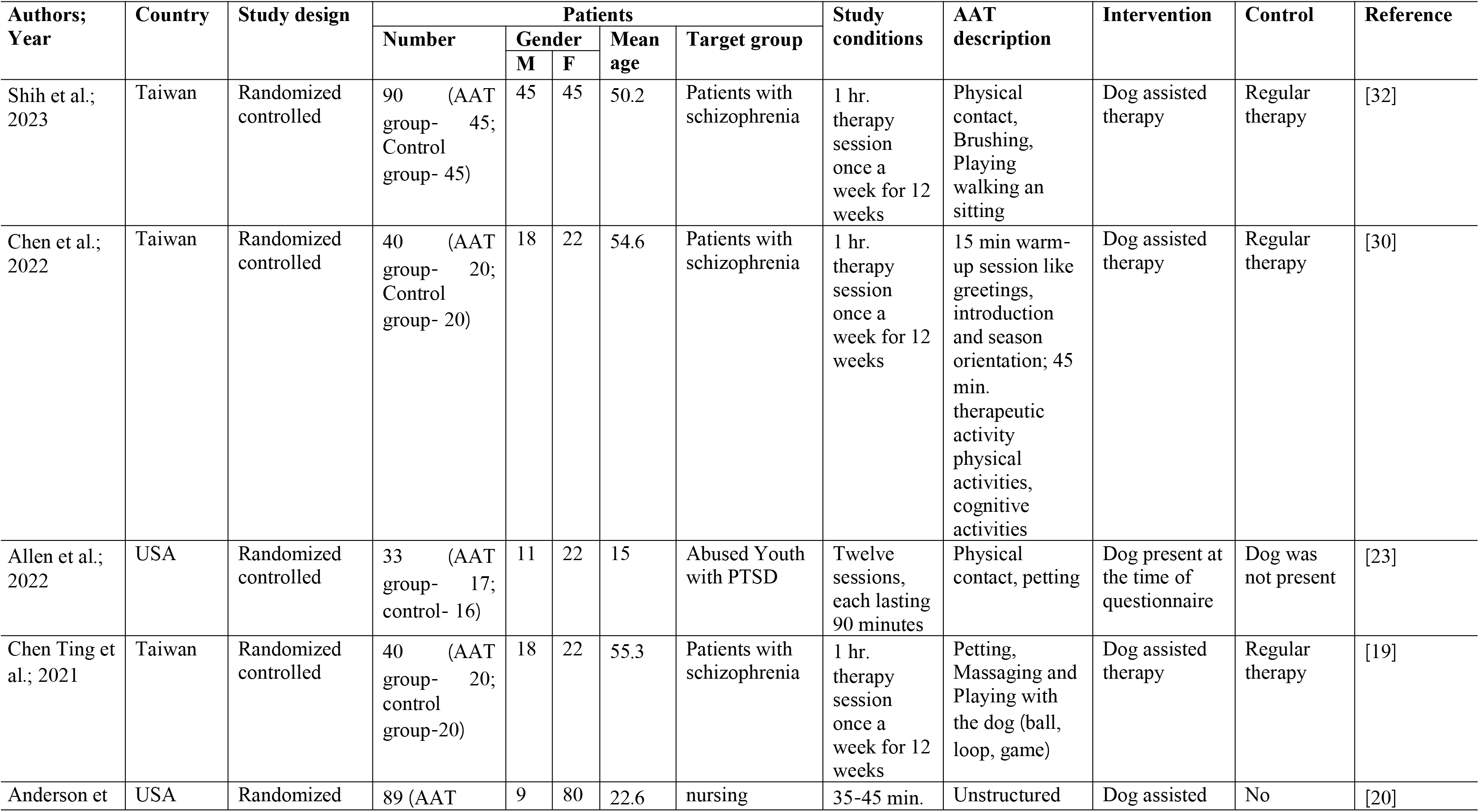
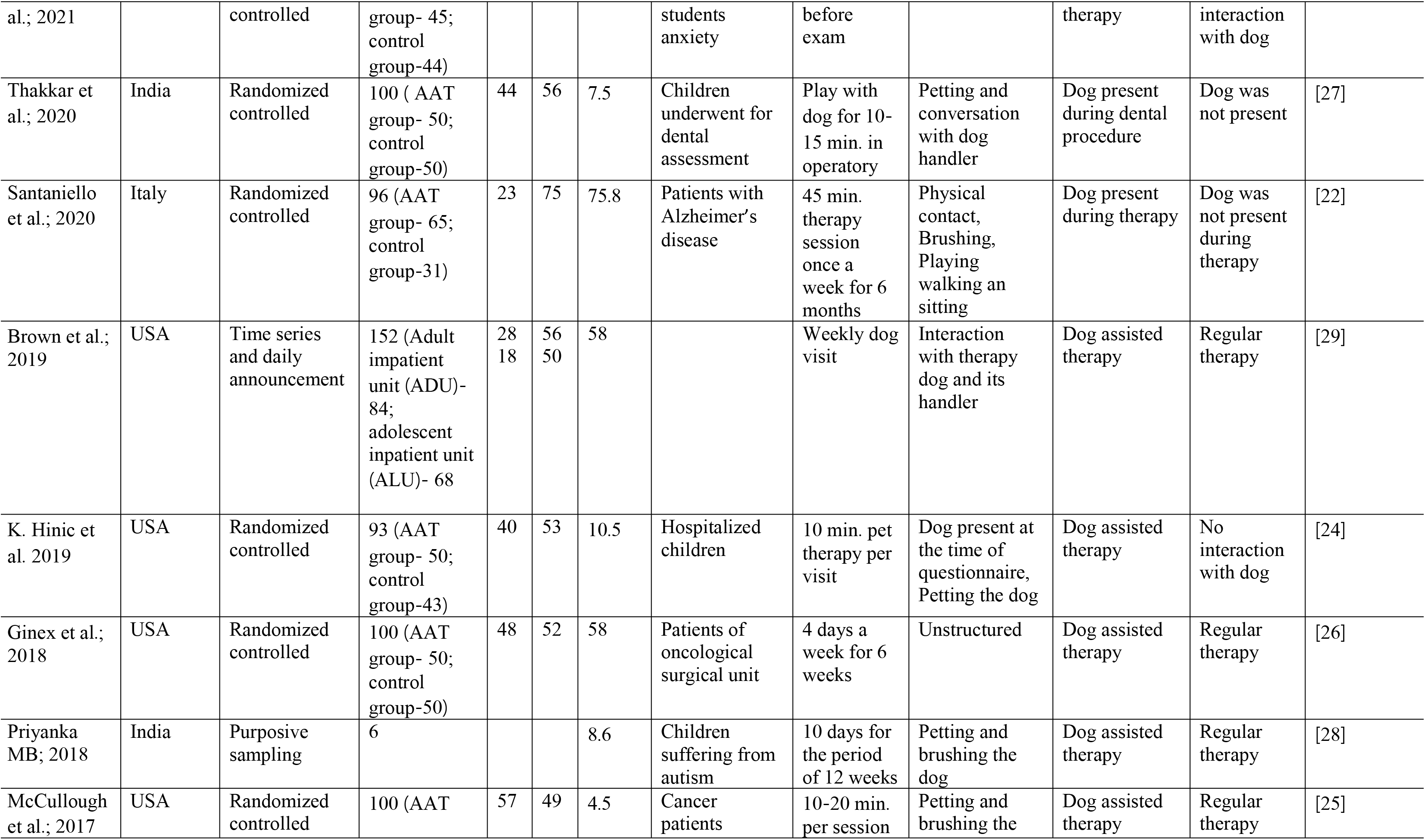
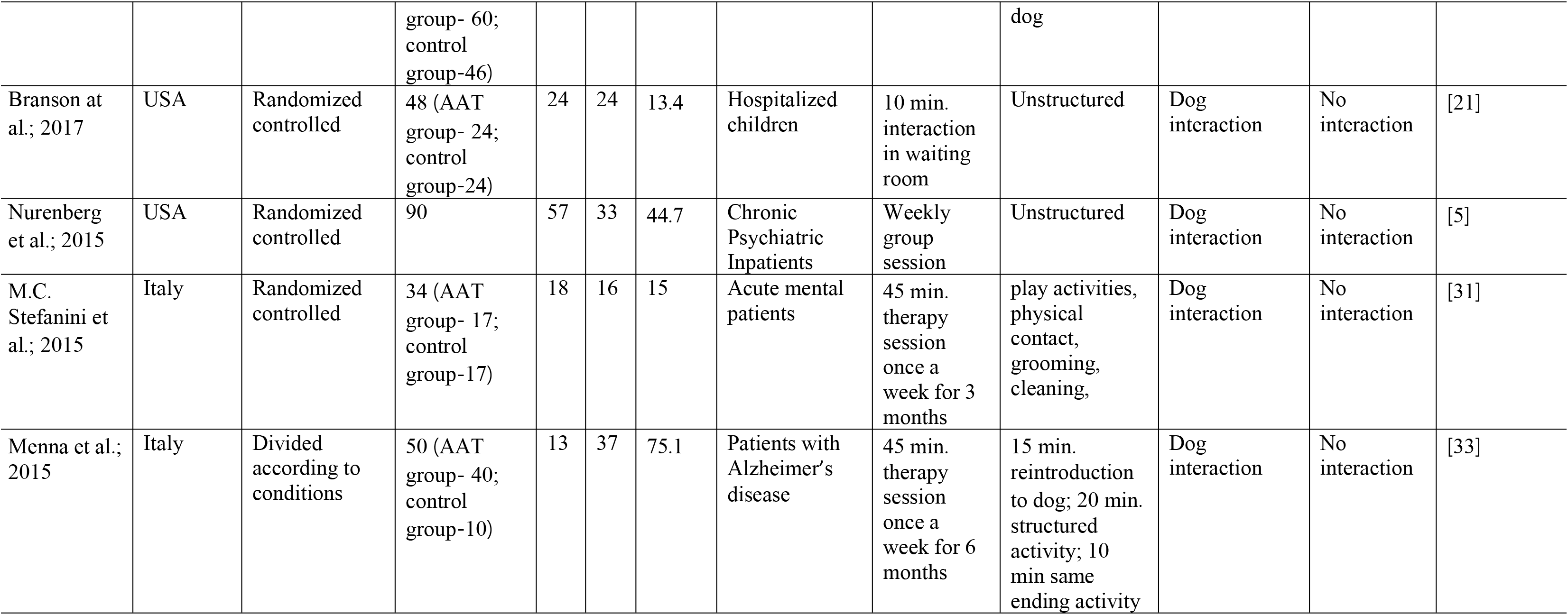
Study characteristics.

**Table 2:** Summary of outcomes from studies included.

| Author; Year | Type of disorder | Mediation | Study Outcome of AAT | Conclusion | Reference |
| --- | --- | --- | --- | --- | --- |
| Shih et al.; 2023 | Psychiatric disorder | Therapeutic | Mental health-social functioning scale (MHSFS); Social adaptive function scale (SAFS); Taiwanese version of the world health organization quality of life (WHOQOL) | Social functioning was significantly higher in the experimental group, Quality of life improved | [32] |
| Chen et al.; 2022 | Psychiatric disorder | Therapeutic | Montreal Cognitive Assessment (MOCA), Chair Stand Test (CST); Timed Up-and-Go (TUG); 5 min walk test; Assessment of Communication and Interaction Skills (ACIS) | Significant improvements in communication and interpersonal skills; improve lower extremity strength and social functions | [30] |
| Allen et al.; 2022 | Stress and anxiety | Activating | Service Satisfaction Scale (SSS); Posttraumatic Stress Disorder Reaction Index for DSM-5; Strengths and Difficulties Questionnaire (SDQ); Moods and Feelings Questionnaire (MFQ); Screen for Child Anxiety Related Disorders (SCARED) | Improvements in caregiver-reported PTSD symptoms, internalizing concerns, and externalizing problems | [23] |
| Chen Ting et al.; 2021 | Psychiatric disorder | Therapeutic | Positive and Negative Syndrome Scale (PANSS), Depression Anxiety Stress Scales Assessment (DASS), and Chinese Happiness Inventory (CHI). | ↓ of stress in the AAT group than in the control group | [19] |
| Anderson et al.; 2021 | Stress and Anxiety | Supportive | State-Trait Anxiety Inventory (STAI) | decreased anxiety in a convenience sample | [20] |
| Thakkar et al.; 2020 | Stress and Anxiety | Activating | Modified Faces Version of the Modified Child Dental Anxiety Scale (MCDASF) | Effective for behavior management strategy for children | [27] |
| Santaniello et al.; 2020 | Neurological disorder | Therapeutic | Mini Mental State Examination (MMSE), Geriatric Depression Scale (GDS) | AAT showed an improvement in both cognitive function and mood | [22] |
| Brown et al.; 2019 | Psychiatric disorder | Therapeutic | Wilcoxon signed-rank test | Positive therapeutic impact on patients and staff in acute care psychiatric units, promoting positive mood and emotions. | [29] |
| K. Hinic et al. 2019 | Stress and Anxiety | Activating | State-Trait Anxiety Scale for Children (STAIC) | Reduce anxiety in hospitalized children and enhance family satisfaction. | [24] |
| Ginex et al.; 2018 | Depression | Supportive | Patient Health Questionnaire-4 (PHQ 4) | Promotes a healing environment for patients that involves a holistic and humanistic perspective | [26] |
| Priyanka MB; 2018 | Neurological disorder | Therapeutic | Autism Treatment Evaluation Checklist (ATEC) and semi-structured interview for 15 min. | Improved expression and communication skills when interacting with the dog, as well as noticeable enhancements in social and motor abilities. | [28] |
| McCullough et al.; 2017 | Stress and anxiety | Supportive | State-Trait Anxiety Inventory (STAI) | Help in reducing the stress and anxiety levels | [25] |
| Branson et al.; 2017 | Stress and anxiety | Supportive | State-Trait Anxiety Inventory for children (STAI-C); Positive and Negative Affect Schedule for Children (PANAS-c) | Increase positive feeling in hospitalized children | [21] |
| Nurenberg et al.; 2015 | Psychiatric disorder | Therapeutic | Equine Assisted Growth and Learning Association (EAGALA); equine-assisted psychotherapy (EAP), canine-assisted psychotherapy (CAP) | Effective therapeutic modality for long-term psychiatric patients at risk of violence | [5] |
| M.C. Stefanini et al.; 2015 | Psychiatric disorder | Supportive | Children Global Assessment Scale (CGAS) | Significant improvements in overall functioning, a decrease in the need for specialized care, and an increase in regular school attendance. | [31] |
| Menna et al.; 2015 | Neurological disorder | Therapeutic | Mini-Mental State Examination (MMSE), 15-item Geriatric Depression Scale (GDS)) | Improve cognition and mood through repeated multimodal stimulation. | [33] |

### 3.2 General characteristics

To provide a concise summary of the selected studies, we have compiled an overview in Table 1. It presents key information from each study, allowing for a quick and comprehensive understanding of the research landscape.

#### 3.2.1 Quantitative study

The studies analyzed in this research were conducted between 2015 and 2023, resulting in a total of sixteen [5, 19–33]included papers. These studies were carried out in various countries, including the USA (n = 8), Taiwan (n = 3), India (n = 2), and Italy (n = 3). Among the included studies, approximately 81.25 percent utilized a randomized controlled trial design. Two studies employed conditional controlled designs, while one study followed a Time series and daily announcement approach. The number of patients enrolled in the studies varied, ranging from 6 to 152 individuals. Specifically, five studies involved 0–40 patients, two studies included 41–80 patients, eight studies comprised 81–120 patients, and one study encompassed 121–160 patients. The selected studies covered a diverse range of populations, with seven studies focusing on children and adolescents and same number of studies involving adult patients, and two studies specifically targeting old people. In the gender distributions, women were more prominently represented, with ≥50% female participants in 13 out of the 16 studies. (Table 1)

The interventions included in the studies were described using various terms such as pet encounter therapy, pet-facilitated therapy, pet-assisted living, animal-assisted intervention, animal-assisted therapy, animal-assisted activity, or simply dog visits/therapy. All of the studies incorporated dogs as the primary intervention. In terms of the duration of the interventions, the seven studies had varying time periods per visit [5, 20, 21, 24, 25, 27, 29]. Five studies had interventions lasting for twelve weeks[19, 28, 30–32] and one for six weeks [26] while two studies had longer intervention periods of six months[22, 33]. One study did not explicitly mention the duration of the intervention [23]. The majority of the studies employed a one-to-one approach in delivering the intervention, emphasizing individual interactions between participants and the dogs. However, one study was conducted in a group setting [5]. Most studies actively encouraged touching and interaction with the animals, while in two studies, the interaction was described as unstructured [20, 21].

#### 3.2.2 Qualitative study

The description of selected studies for qualitative analysis is presented in Table 1 and 2. Studies reported on the experience of the patients with animal assisted intervention such as dog therapy or animal visit. Thematic analysis was employed to identify recurring themes and extract meaningful insights from the type of disorder. Participants described the animals as a source of comfort, providing emotional support and reducing stress and anxiety. The interactions with the animals were reported to have a soothing effect and helped individuals cope with their challenges and emotional difficulties.

The qualitative analysis shed light on the subjective experiences and perceptions of individuals participating in the interventions. It provided valuable insights into the emotional, social, and therapeutic benefits associated with animal-assisted interventions, highlighting the potential of these interventions to enhance well-being and quality of life. The selected studies pertaining to psychiatric disorders predominantly focused on schizophrenia, with five studies specifically addressing this condition in adults. Additionally, one study explored acute mental disorders in children [31]. Six studies were dedicated to investigating stress and anxiety, targeting various populations such as children undergoing physical examinations, children with post-traumatic stress disorder (PTSD), patients with cancer, and nursing students. Three studies examined neurological disorders, including one study involving children with autism[28] and two studies involving older individuals with Alzheimer’s disease[22, 33]. In one study, the intervention aimed to reduce depression among patients undergoing oncology surgery [27].

### 3.3 Outcomes

The number of studies with at least one statistically significant positive outcome measure, divided by patient condition and intervention category, is presented in Table 3. The study aimed to comprehensively evaluate mental health, social functioning, and overall quality of life, taking into account various parameters specific to each measurement scale e.g. generic health related quality of life measures like, Posttraumatic Stress Disorder Reaction Index for DSM-5, State- Trait Anxiety Scale for Children (STAIC), Patient Health Questionnaire-4 (PHQ 4), Patient Health Questionnaire-4 (PHQ 4), Mini-Mental State Examination (MMSE), 15-item Geriatric Depression Scale (GDS). General functional measure e.g. mental health-social functioning scale (MHSFS), Social adaptive function scale (SAFS), Chair Stand Test (CST), Timed Up-and-Go (TUG), Assessment of Communication and Interaction Skills (ACIS), etc.

**Table 3:** Number of studies classified based on the condition, type of intervention, and the presence of positive outcome.

| Condition | Supportive mediation |  | Therapeutic mediation |  | Activating mediation |  |
| --- | --- | --- | --- | --- | --- | --- |
|  | Yes | No | Yes | No | Yes | No |
| Psychiatric disorder | 1 | - | 4 | 1 | - | - |
| Neurological disorder | - | - | 3 | - | - | - |
| Stress and anxiety | 3 | - | - | - | 2 | 1 |
| Depression | 1 | - | - | - | - | - |

#### 3.3.1 Psychiatric disorder

All six trials that focused on psychiatric disorders were categorized as AAT and involved interventions with dog therapy. Among these studies, five were conducted using a randomized controlled design, while one study utilized a time series design with randomized daily announcements within a pre-post experimental framework. One study specifically examined patients in child and adolescent psychiatry [31], while the remaining five studies focused on adult psychiatry patients [5, 19, 29, 30, 32]. The duration of the AAT programs varied, with some studies consisting of 12-week programs in different settings, while two studies provided weekly therapy sessions without specifying the intervention period [5, 29].

Each of the six conducted studies involved a comparison between an intervention group receiving a specific therapy and a control group that did not participate in any related activities. Notably, the five studies specifically targeted middle-aged and older patients diagnosed with chronic schizophrenia. The results of these studies consistently demonstrated significant improvements in various areas, including reductions in psychiatric symptoms, enhanced social functioning, improved quality of life, enhanced cognitive function, increased agility and mobility, and decreased stress levels. These outcomes were measured using a variety of scales and assessment tools[5, 19, 30–32]. In a study conducted by Brown et al.[29], the focus was on examining the impact of mood states and feelings among patients and staff in inpatient psychiatric units. The researchers observed significant changes in mood before and after sessions involving therapy dogs. Specifically, negative moods decreased, while positive moods, such as feelings of happiness, relaxation, and calmness, increased. These changes were measured using the Visual Analog Mood Scale[29]. Overall, these findings highlight the efficacy of AAT in positively impacting the well-being and overall functioning of individuals with psychiatric disorder.

#### 3.3.2 Neurological disorder

Among the studies that focused on neurological disorders, three of them utilized dog therapy as an intervention. One of these studies employed a randomized control design [22], while the other two studies used purposive sampling based on the patients’ condition. One study specifically targeted children and adolescents with autism, while the other two studies focused on elderly patients with Alzheimer’s disease. The duration of the AAT programs varied, ranging from 3 to 6 months.

In each of the three conducted studies, the intervention group was compared to a control group that did not participate in any activities, in order to assess the outcomes of the therapy. Priyanka MB’s[28] study focused on children with autism and observed that engaging with a therapy dog, such as brushing the dog and attempting to draw and write for the dog, led to enhanced social and motor skills. Additionally, the children experienced a sense of relaxation and calmness in the presence of the dog. The studies conducted by Menna et al.[33] and Santaniello et al.[22], focusing on elderly patients with Alzheimer’s disease over a period of six months, have shown promising results. Menna et al.’s[33] study demonstrated the applicability and effectiveness of AAT interventions in stimulating cognition and improving mood. The interventions involved repeated multimodal stimulation, including verbal, visual, and tactile approaches. Similarly, Santaniello et al.’s[22] study also revealed improvements in both cognitive function and mood in the AAT group, as measured by changes in the MMSE and GDS. Overall, these studies indicate that non-pharmacological therapies, particularly AAT, have the potential to reduce symptoms associated with neurological disorders.

#### 3.3.3 Stress and anxiety

The six trials that specifically addressed stress and anxiety utilized AAT interventions involving dog therapy. These studies exclusively targeted children and adolescents, employing a randomized controlled design. In five of the studies, the therapy sessions lasted between 10 to 45 minutes, while one study did not specify the duration of the intervention period.

The study conducted by Allen et al.[23] focused on youths who had experienced abuse and were diagnosed with posttraumatic stress disorder (PTSD). The results revealed that the group receiving the intervention showed greater improvements in caregiver-reported symptoms of PTSD, internalizing concerns, and externalizing problems compared to the control group[23]. In a study by Anderson et al.[20] involving nursing students, the intervention group experienced interactions with dogs before testing. This interaction served as a stress reliever for the students, resulting in a decrease in anxiety as measured by the STAI. Thakkar et al.[27] conducted a study on children who were undergoing dental assessments. The findings indicated that the intervention group showed a significantly greater reduction in anxiety compared to the control group, as measured by the Modified Faces Version of the modified child dental anxiety scale (MCDASF). In the studies conducted by K. Hinic et al.[24] and Branson et al.[21], dog therapy was provided to hospitalized children, and their anxiety levels were assessed before and after the intervention. The results from the STAI-C suggested that brief pet therapy visits served as a tool to decrease anxiety in hospitalized children and promote family satisfaction. McCullough et al.[25] conducted a study where the intervention group participated in dog therapy, while the control group received standard care at the hospital. The findings demonstrated the applicability and effectiveness of animal-assisted therapy (AAT) interventions in reducing stress and anxiety levels in cancer patients.

Overall, when considering the results of all these studies, it becomes evident that each one exhibited at least one statistically significant positive effect. When these findings are examined collectively, they provide compelling evidence to suggest that particular modalities of AAT hold substantial promise in terms of reducing stress levels and fostering a positive impact on individuals’ overall mood and well-being.

#### 3.3.4 Depression

Ginex et al.[26] conducted a study to explore the impact of dog-assisted intervention on an inpatient surgical oncology unit. The study employed a randomized control design, with patients in the intervention group receiving therapy four days a week throughout the study period. In contrast, the control group underwent physical therapy without any modifications to their normal routine. Patients in the intervention group reported a significant decrease in depression and anxiety levels, as measured by the Patient Health Questionnaire-4 (PHQ-4), compared to the control group. The findings of the study suggest that AAT fosters a healing environment for patients, incorporating a holistic and humanistic approach that elicits overwhelmingly positive responses.

## 3. Discussion

The outcomes of this meta-analysis provide the longstanding belief that animals can play a beneficial role in the healing process. The study revealed positive and moderately strong results across various aspects, including medical well-being, behavioral outcomes, and the reduction of Autism spectrum symptoms. Moreover, the effect sizes in all four outcome areas were consistent and uniform. Additionally, support for AAT was evident from four studies comparing it with established interventions, demonstrating that AAT was equally or more effective. These compelling findings indicate that AAT is a robust intervention deserving of further exploration and utilization. This systematic review and meta-analysis specifically focused on dogs as the assisting animals in a healthcare setting. However, there were no limitations on the characteristics of the population included in the study. Although this research synthesis provides evidence in favor of the effectiveness of AAT, it is essential to acknowledge the complexities associated with interventions in general and the specific nuances related to the utilization of AAT.

The majority of articles included in this systematic review were based on randomized controlled trials conducted in various countries. Additionally, time series and daily announcements, divided according to different conditions, were also considered. The increased number of studies provided greater power in assessing variance heterogeneity and potential group differences.

Although the results are speculative, the meta-analysis demonstrated homogeneity in the summary values, with only one exploratory group difference reaching statistical significance. Nonetheless, this analysis brought forth several intriguing questions and patterns, serving as a foundation for discussions or further research on the factors influencing the effectiveness of AAT. For instance, consistent benefits were observed in children, young age groups, and old age groups across all outcome variables, including symptoms associated with psychiatric disorders, stress, and anxiety. In particular, among the adult group, a high prevalence of psychiatric disorders, followed by neurological disorders, stress, and anxiety, was found. In contrast, in children, a high number of cases related to stress and anxiety disorders were identified.

Several organizations in different countries are actively working to promote animal-assisted therapy. In the United States, The Society for Healthcare Epidemiology of America (SHEA) has established comprehensive guidelines for animals in healthcare facilities, which emphasize the importance of written policies, designated AAI visit liaisons, and formal training programs for animals and handlers[34]. However, despite these guidelines, there is no legal requirement for healthcare facilities to adopt these measures. One notable organization in the US, Pet Partners, stands out as the only national therapy animal organization that mandates volunteer training, biennial evaluations of animal-handler teams, and prohibits raw meat diets [35]. In Europe, the European Society for Animal Assisted Therapy (ESAAT) plays a significant role as an influential organization operating across various disciplines and professions within the field. ESAAT’s primary mission is to accredit education and training programs in the domain of animal-assisted interventions[36]. While the Western world has made significant advancements in animal-assisted therapy, Eastern countries such as India, China, Taiwan, Japan, and Sri Lanka are still in the early stages of exploring and implementing such practices. These countries are currently in the infancy phase of utilizing and developing their own animal-assisted therapy programs. As awareness and understanding of the benefits of animal-assisted therapy continue to grow globally, it is expected that these Eastern countries will gradually catch up and further enhance their animal-assisted therapy initiatives[37].

Our review was based on a limited number of studies, it can be attributed due to our strict inclusion criteria and the presence of suboptimal study designs. Specifically, many of the randomized trials were characterized by small sample sizes, short durations, and a lack of follow-up assessments. Additionally, some trials suffered from a lack of blinding, as both the participants and researchers were aware of the group allocation, which may introduce bias. Another limitation pertains to the suitability of the outcome measures used, which may not fully capture the important values and impacts as perceived by the participants. On the other hand, the qualitative research included in the review exhibited higher overall quality and contributed valuable insights to our findings.

## 4. Conclusion

In conclusion, the reviewed studies provide preliminary evidence of the potential benefits of AAT in certain conditions. It suggest that dog-assisted therapy can have minor to moderate effects in treating psychiatric disorders, cognitive disorders, neurological disorders etc., and demonstrates potential in various medical interventions. However, it is important to note that some of outcome measures analyzed did not show significant effects and further research is needed to better understand the specific contexts and conditions. To foster growth of such therapy, we need education campaigns, research programs, professional support, and media awareness to increase the effectiveness of AAT across different countries.

## Ethics approval and consent to participate

Not Applicable

## Consent for publication

Not Applicable

## Availability of data and material

The authors confirm that the data supporting the findings of this study are available within the article

## Author’s Contributions

All authors contributed to writing-original draft preparation, review and editing.

## Acknowledgements

We greatly appreciate VtR Inc-CGU (SCRPD1L0221); DOXABIO-CGU (SCRPD1K0131), and CGU grant (UZRPD1L0011, UZRPD1M0081).

